# Graph neural network approach for inferring neuronal connectivity

**DOI:** 10.64898/2026.10.28.762597

**Authors:** Archit Gupta, Curtis L. Neveu, John H. Byrne

## Abstract

Understanding the connectivity of neuronal networks is essential for understanding brain function. Current anatomical mapping techniques, such as serial section microscopy and viral tracing, are resource-intensive and cannot capture the efficacy of connectivity. A complementary approach is to infer effective connectivity from single-neuron-resolution population activity. We developed a graph neural network (GNN)-based tool, CircuitGNN, that accurately infers neural effective connectivity from population activity. CircuitGNN accurately inferred the connectivity of several small circuits and detected both the strengthening and weakening of connections, such as those that occur during learning. CircuitGNN outperformed other methods when applied to simulated population activity generated by an invertebrate central pattern-generating circuit and a cortical microcircuit. CircuitGNN was evaluated on networks of varying size, recording duration, and proportion of hidden neurons. Although performance improved with longer recordings, CircuitGNN achieved high precision with recordings as short as 4 min, scaled to large circuits of up to ~750 neurons, and remained robust when 20% or 40% of neurons were hidden. These results establish CircuitGNN as an accurate and robust technique for inferring neural connectivity from activity.

## INTRODUCTION

Understanding the connectivity of neuronal networks is essential for understanding brain function. Although mapping of neuronal circuits by traditional methods such as serial section electron microscopy [1–3] and viral tracing [4] has been successful for some organisms, these methods remain tedious and impractical, and they cannot capture the efficacy of connections. Importantly, these methods provide only a snapshot of the connectivity and are limited in detecting changes in efficacy associated with learning, especially when learning involves changes in intrinsic electrical excitability of neurons [5]. New technologies can simultaneously record from thousands of neurons. Several approaches have been developed to infer connectivity from these large datasets. These connectivity inference techniques rely on cross-correlation [6,7], information theory [8–11], generalized linear models [12–14], machine learning [15], or deep learning [16–19]. These techniques often struggle to identify inhibitory and weak connections [9], are confounded by input from unobserved neurons, and scale poorly to large populations [20]. Some techniques ignore inhibitory-to-inhibitory connections entirely [6], a significant limitation given that these connections constitute a substantial component of local cortical circuits [21].

Graph neural networks (GNNs) are deep learning models that use data to extract representations from graphs, which can then be used to answer questions about the graph’s nodes, edges, or the full graph [22,23]. These representations are high-dimensional vectors that encode task-relevant information in a machine-understandable format [23]. Formally, a graph *G* = (*V, E*) consists of nodes *V* connected by edges *E*. The core component of the GNN is the message-passing layer, in which each node collects messages in the form of vector representations from its neighboring nodes, aggregates them into a single vector, and uses this vector to update its own representation [23]. Through this process, each node can iteratively capture increasingly complex patterns from its surrounding graph structure. After the final message-passing layer, these representations are used to make predictions about the properties of nodes, edges, or the entire graph. Although several GNN-based methods have been previously developed [24,25], they cannot distinguish between excitatory and inhibitory connections, have been evaluated only in limited settings (e.g., aggregated population activity [24] or ring attractor networks [25]), and lack easy-to-use implementations, limiting their widespread adoption.

Here, Circuit Graph Neural Network (CircuitGNN) is introduced to accurately infer neural connectivity. CircuitGNN represents connectivity using two nonnegative adjacency matrices, corresponding to excitatory and inhibitory strength, which are combined into a single signed connectivity matrix by taking their difference. This signed connectivity matrix is used across successive rounds of message passing, allowing the model to capture nonlinear, polysynaptic interactions. CircuitGNN is trained in a self-supervised manner to predict neural activity at the next time step from recent history, requiring no prior knowledge of connectivity and making it directly applicable to large-scale neural recordings.

## RESULTS

### Circuit Graph Neural Network

The goal of inferring connectivity between neurons is a graph structure learning (GSL) problem, in which a graph is learned through the optimization of a downstream task [26–28]. CircuitGNN is a GSL framework comprising a structure learner that learns the graph structure, and a GNN that uses the learned structure to optimize the downstream task. Here, the downstream task is prediction of future activity from past activity of the entire population (Fig. 1A). The framework comprises two modules. The first is a spike encoder, which learns a low-dimensional representation of each neuron’s activity (Fig. 1B). The second is a graph neural network (GNN) whose core operation is message-passing (Fig. 1C). Connectivity is stored in an adjacency matrix, initialized randomly at the start of training; this matrix is the structure learner. Each entry describes the connection between a pair of neurons. An entry’s sign specifies whether the connection is excitatory or inhibitory and its magnitude specifies the strength. This matrix can equivalently be viewed as a graph (Fig. 1C1). At each message-passing layer, every neuron updates its representation by aggregating information from all other neurons (Fig. 1C2). For a given target neuron (e.g., neuron A in Fig. 1), this incoming information is combined through three terms. In the additive term, every other neuron’s representation is scaled by its connection weight to the target neuron and summed. In the multiplicative term, every other neuron’s representation is scaled by both its connection weight to the target neuron and the target neuron’s own representation, then summed. The self term is a transformation of the target neuron’s own representation. In this example, neurons B and D are randomly initialized as connected to neuron A, so only their representations contribute to updating the representation of neuron A in the first iteration (Fig. 1C2). Because the initial connectivity is arbitrary, it is refined as training proceeds. The additive, multiplicative, and self terms are combined to produce each neuron’s updated representation (Fig. 1C3), which is then used to predict whether the neuron will spike in the future (Fig. 1D). This prediction is compared to the recorded activity (Fig. 1E), the resulting error is backpropagated through the model, and gradient descent iteratively updates both the GNN parameters and the inferred connectivity structure. Over many iterations, this optimization yields a learned adjacency matrix (Fig. 1F) that is interpreted as connectivity (Fig. 1G). Because accurate forecasting requires capturing the dependencies that drive population activity, the learned adjacency matrix reflects the connectivity structure of the underlying circuit. Further details are provided in the Methods.

**Figure 1.**
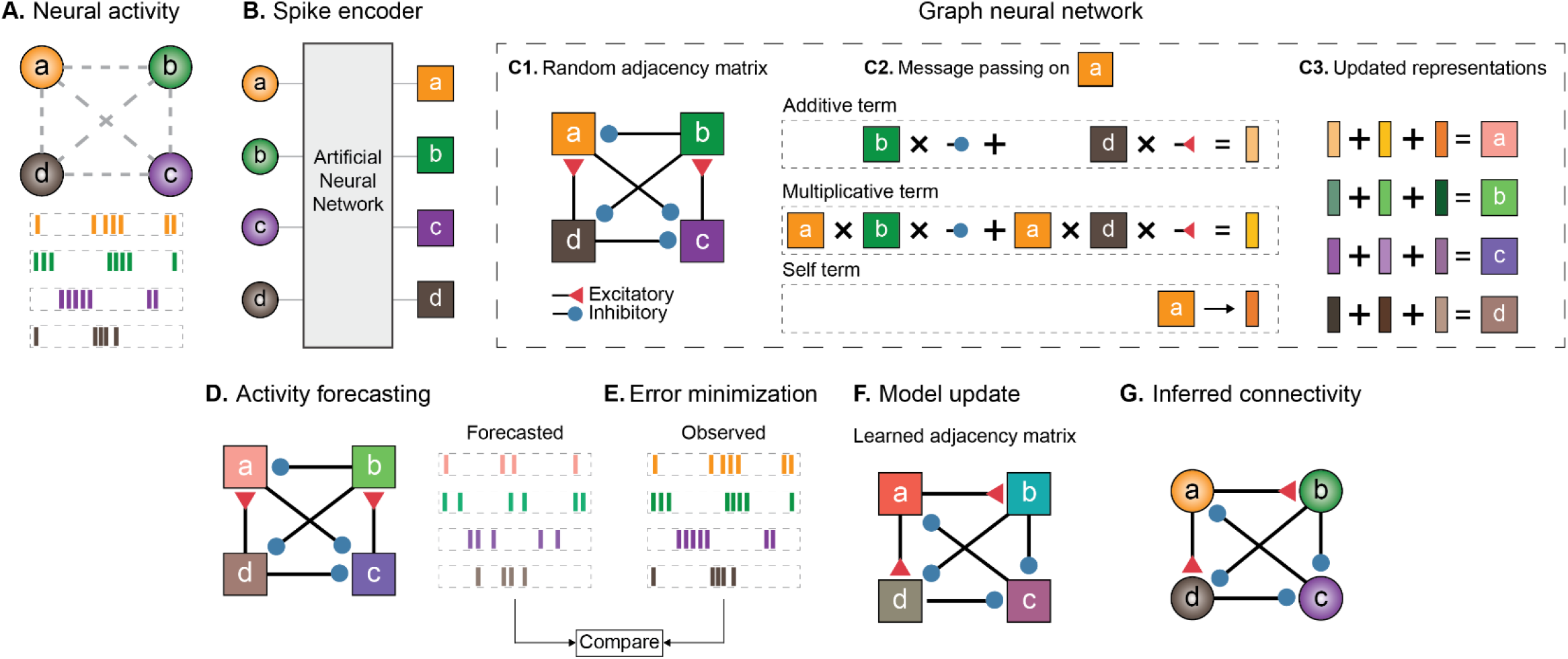
CircuitGNN infers neural connectivity via message passing. **A**. Recorded neural activity from a circuit of four neurons whose connectivity is unknown. **B**. A spike encoder transforms each neuron’s activity into a low-dimensional representation for use by the graph neural network. **C1**. At the start of training, a random adjacency matrix is initialized, representing the connections between neurons; red triangles denote excitatory connections and blue circles denote inhibitory connections. **C2**. Message passing occurs at every neuron, and is illustrated here for neuron A. Three terms are obtained for each neuron: additive and multiplicative terms, which incorporate information from other neurons, and a self term, which incorporates only its own information. **C3**. The three terms are combined into an updated representation for each neuron. **D**. The updated representations are used to predict each neuron’s future activity. **E**. Predicted and recorded activity are compared, and the error is minimized. **F**. Over many iterations, the graph neural network learns an adjacency matrix. **G**. The learned adjacency matrix is interpreted as the inferred connectivity.

### Validation of CircuitGNN on simulated data

To explore CircuitGNN’s ability to capture neural connectivity, CircuitGNN was applied to small simulated circuits that have previously been used to validate connectivity inference algorithms [9,10]. Specifically, three conductance-based neuronal models were examined, implemented via Simulator for Neural Networks and Action Potentials (SNNAP), which simulates neurons using Hodgkin-Huxley-type conductance-based equations along with various chemical and electrical synapses [29]. Networks A (Fig. 2A1) and B (Fig. 2B1) consisted of three neurons, in which neuron A was activated by a depolarizing current (0.57 nA, 10 s) delivered at an interstimulus interval of 10 s. Network A included an excitatory and an inhibitory synapse, whereas network B included three excitatory synapses. CircuitGNN successfully inferred the connectivity of both networks (Fig. 2A2, 2B2). Network C (Fig. 2C1) consisted of 11 neurons and included four motifs: an extended path of serially connected nodes (nodes 1, 4, 11), a sink node influenced by multiple nodes (node 2), a multipath connection from node 6 to node 10, and an isolated node (node 7). True connectivity is shown in Fig. 2C2, and connectivity inferred by CircuitGNN is shown in Fig. 2C3. As with networks A and B, CircuitGNN successfully inferred the connectivity of network C without errors.

**Figure 2.**
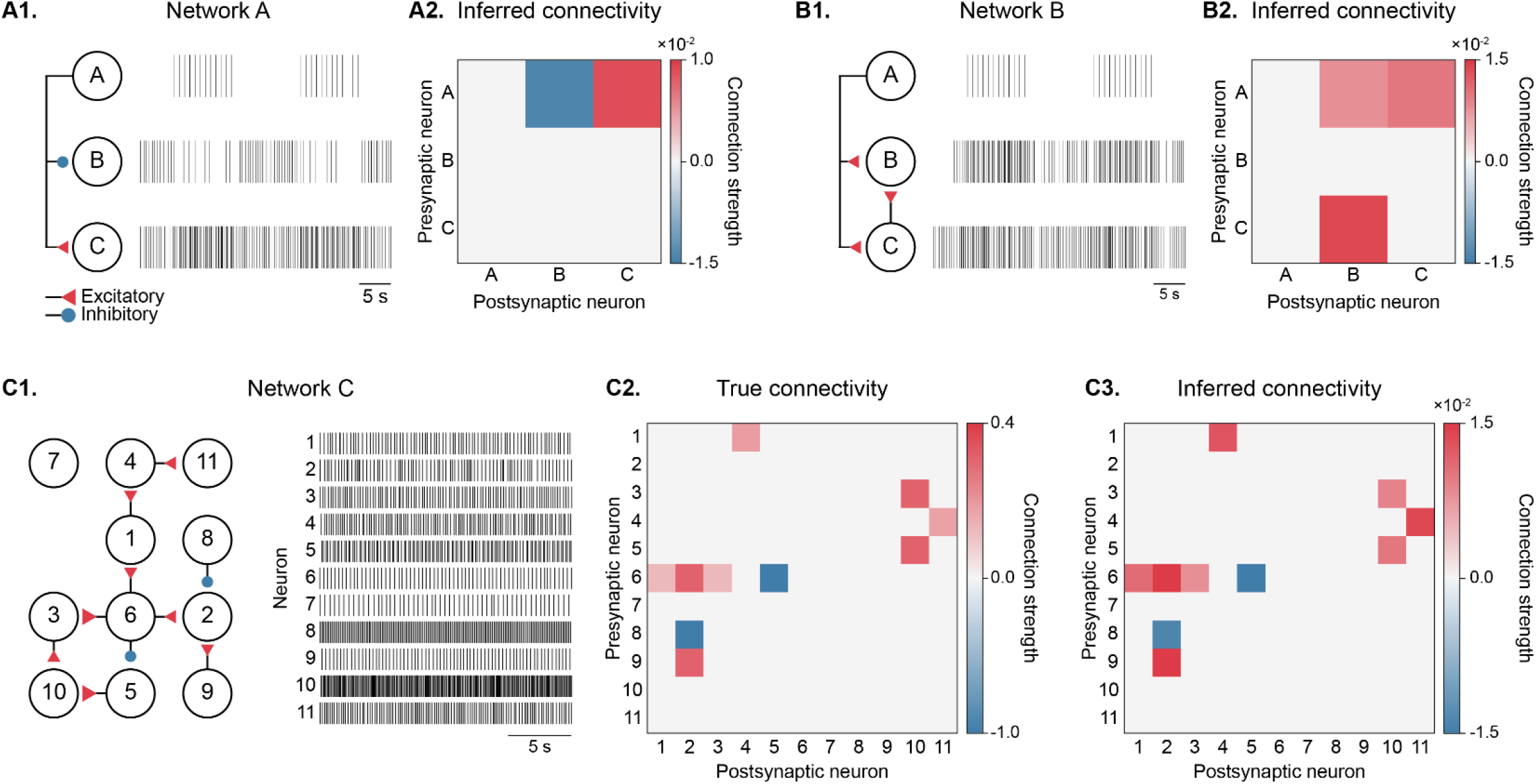
Validating CircuitGNN on simulated data. **A1**. Network connectivity and spiking activity of network A. Red triangles denote excitatory connections; blue circles denote inhibitory connections. **A2**. Network connectivity inferred by CircuitGNN. Positive connections (red) are excitatory; negative connections (blue) are inhibitory. **B1**. Network connectivity and spiking activity of network B. **B2**. Network connectivity inferred by CircuitGNN. **C1**. Network connectivity and spiking activity of network C. **C2**. True connectivity of the network. Connection strengths are derived from conductance values and normalized to the maximum. **C3**. Network connectivity inferred by CircuitGNN.

A useful connectivity inference technique should be able to detect subtle changes in the connectivity of an underlying circuit, such as changes to a single connection’s strength. To test whether CircuitGNN can detect such changes, two variants of network C were developed (Fig. 3). In the first, the connection between neuron 3 and neuron 10 was weakened (Fig. 3B); in the second, the connection between neuron 6 and neuron 2 was strengthened (Fig. 3C). CircuitGNN successfully inferred the weakened connection between neurons 3 and 10: whereas the 3-to-10 and 5-to-10 connections were inferred at comparable weights in the baseline condition, the inferred weight of the 3-to-10 connection fell to 0.46 of the 5-to-10 connection in the first variant, close to the ground truth ratio of 0.40 (Fig. 3B). CircuitGNN also detected the strengthened connection between neurons 6 and 2 (Fig. 3C). Taken together, these results demonstrate that CircuitGNN can successfully identify changes in connectivity such as might occur during plasticity of neural networks during learning.

**Figure 3.**
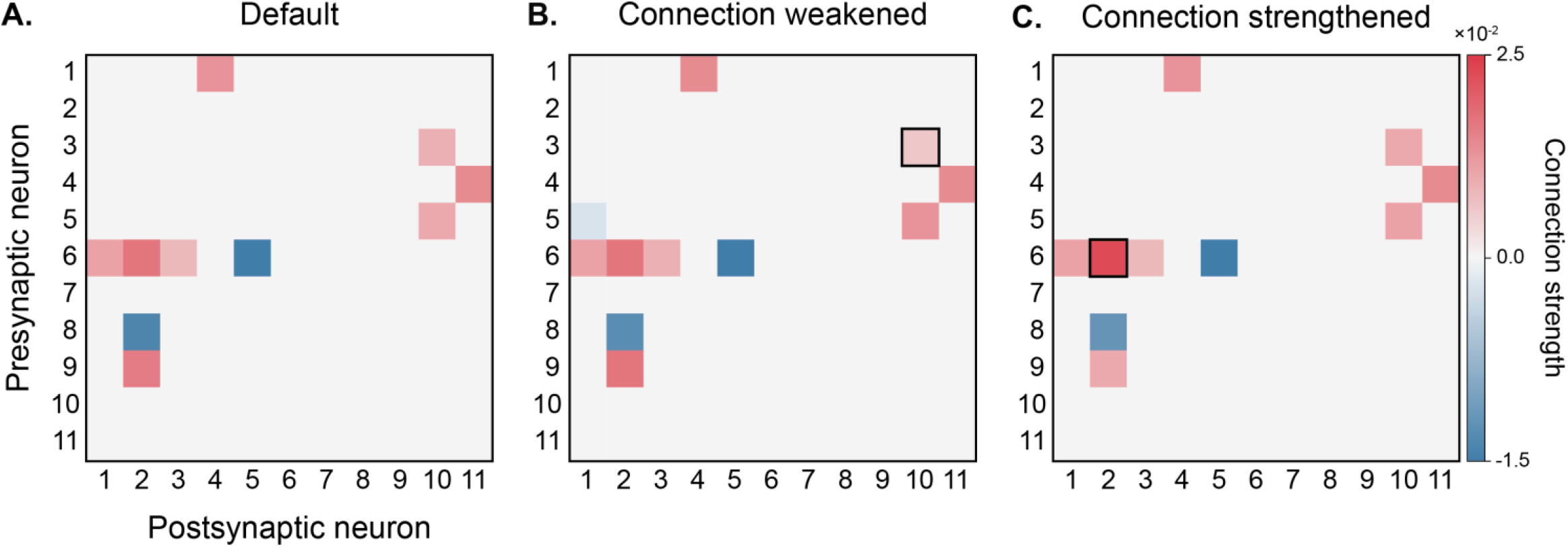
CircuitGNN detects changes to network connectivity. **A**. Connectivity inferred by CircuitGNN under the default condition from Fig. 2C3. **B**. CircuitGNN detects weakened connectivity from neuron 3 to neuron 10 (black square) relative to the connection strength from neuron 5 to neuron 10; synaptic conductance for the neuron 3 to neuron 10 connection was reduced from 1.25 µS to 0.5 µS (60% decrease). **C**. CircuitGNN detects the strengthened connection from neuron 6 to neuron 2 (black square); synaptic conductance for this connection was increased from 1.25 µS to 3.125 µS (150% increase).

### CircuitGNN inferred the connectivity of the feeding circuit of *Aplysia* more accurately than existing methods

CircuitGNN was next applied to population activity generated by a conductance-based computational model of the central pattern generating (CPG) circuit that mediates feeding behavior of the marine mollusc *Aplysia californica* (for reviews, see [29–32]). Inferring the circuit’s connectivity from activity has been challenging for several reasons. First, the activity of the feeding CPG is highly phasic, with each cycle of feeding behavior consisting of two phases and most neurons preferentially active in one or the other. Such phasic activity can cause correlation to be mistaken for connectivity. Second, a single neuron can form both excitatory and inhibitory synapses with its targets. Consequently, connectivity inference cannot be simplified to assigning a single effective sign to each neuron. Third, the feeding network produces multiple types of behaviors (ingestion and egestion), which correspond to distinct patterns of activity.

To evaluate CircuitGNN on the feeding circuit, the conductance-based computational model was used to generate simulated data (Figs. 4A, 5A). Ground truth connectivity is shown in Fig. 4B for a simulation in which operant conditioning, an associative learning paradigm, was modeled [31]. Reconstruction performance was evaluated using F1 score [33] and normalized inference error (NIE) [34]. The F1 score is the harmonic mean of precision and recall, where precision is the fraction of inferred edges that are correct and recall is the fraction of ground truth edges that were inferred. A precision score of 1.0, therefore, means that every edge inferred by CircuitGNN exists in the ground truth, whereas a recall score of 1.0 means that CircuitGNN inferred every edge present in the ground truth. The F1 score thus measures how well CircuitGNN’s inferred edges match the ground truth, with a score of 1.0 indicating that CircuitGNN inferred every ground truth edge without inferring any false edges. NIE measures how much the inferred connectivity matrix differs from the ground truth connectivity matrix. An NIE of 0.0 means that CircuitGNN inferred the ground truth matrix exactly.

**Figure 4.**
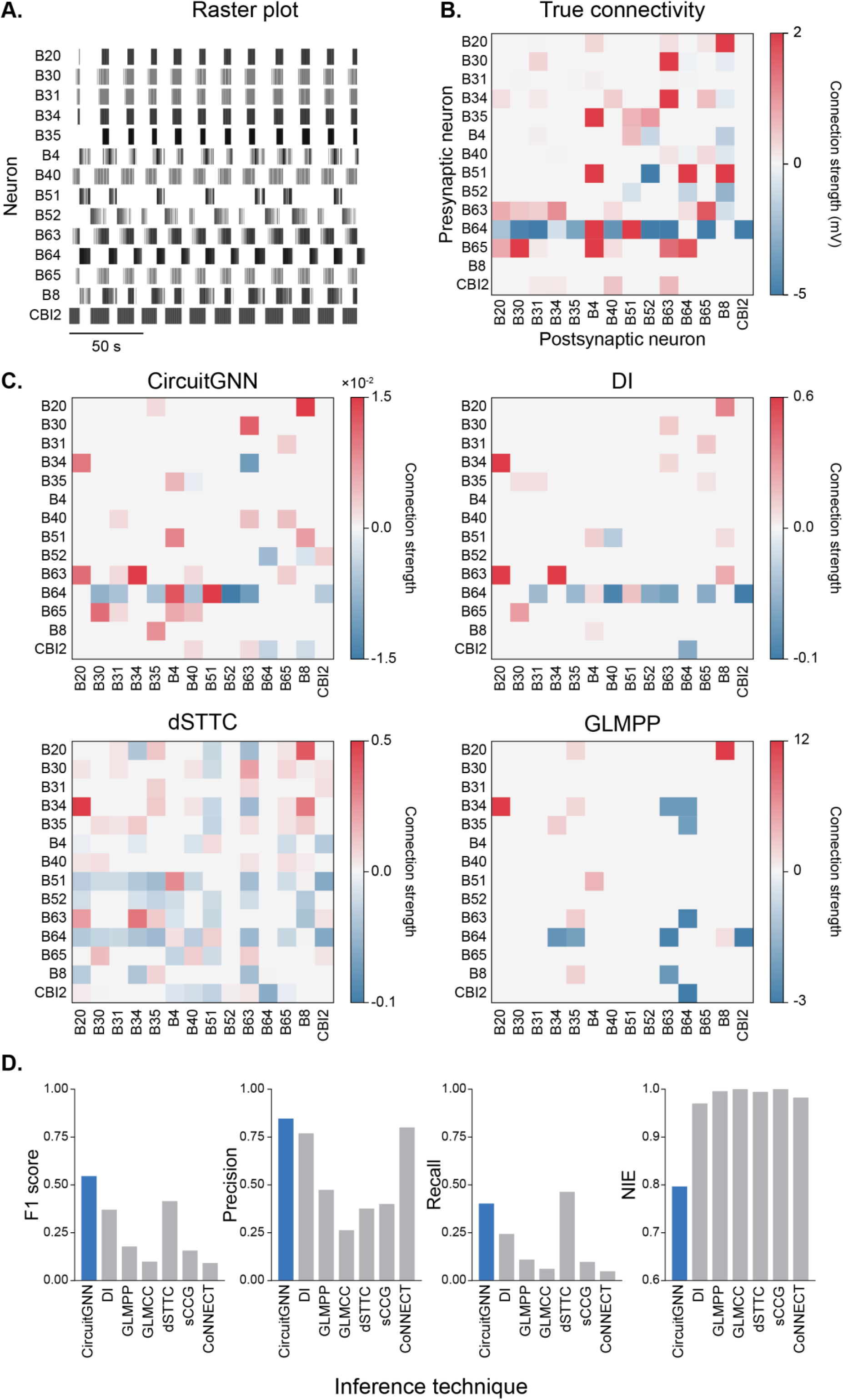
CircuitGNN inferred *Aplysia* feeding circuit connectivity. **A**. Spike raster of the 14 simulated neurons of the feeding central pattern generator during operant conditioning; note the phasic, alternating activity. **B**. Ground truth connectivity in mV. **C**. Connectivity inferred by CircuitGNN and three of the other techniques. **D**. F1 score, precision, recall, and normalized inference error for all evaluated techniques: CircuitGNN, directed information (DI), generalized linear model for point-processes (GLMPP), generalized linear model cross-correlogram (GLMCC), directed spike time tiling coefficient (dSTTC), smoothed cross-correlogram (sCCG), and convolutional neural network for estimating synaptic connectivity from spike trains (CoNNECT).

CircuitGNN was compared against six established connectivity inference techniques: smoothed cross-correlogram (sCCG) [35], directed spike time tiling coefficient (dSTTC) [36], generalized linear model for point-processes (GLMPP) [17,37], generalized linear model cross-correlogram (GLMCC) [13], directed information (DI) [9], and convolutional neural network for estimating synaptic connectivity from spike trains (CoNNECT) [16]. These methods were selected because they are widely used, span a broad range of approaches, and have readily available implementations. To ensure a fair comparison, technique parameters were tuned where applicable, and the best-performing setting for each network was reported (see Methods). CircuitGNN outperformed all other inference techniques, obtaining the highest F1 score (more than 0.13 higher than any other technique) and the lowest NIE (more than 0.15 lower) (Fig. 4C, D, SFig. 1). CircuitGNN obtained a precision of 0.85 and a recall of 0.40, compared with 0.77 and 0.24 for DI, 0.38 and 0.46 for dSTTC, and 0.47 and 0.11 for GLMPP (Fig. 4D). Because any technique can trade recall for precision by retaining only its most confident predictions, differences in precision may partly reflect how conservatively each technique predicts connections. To compare techniques at an equal number of inferred connections, precision@K was also computed, defined as the precision of the K connections with the greatest inferred strength. CircuitGNN sustained high precision over a larger number of connections than other techniques (SFig. 2A).

CircuitGNN identified several connections essential to circuit function that other inference techniques missed (Fig. 4C). Among the excitatory connections, CircuitGNN recovered the connection from the behavior-initiating neuron B63 to another behavior-initiating neuron, B31, which is required for generating feeding motor behavior [38]. CircuitGNN also identified an excitatory connection from CBI-2 to B63, which appears essential for the generation of feeding behavior [38]. Among the inhibitory connections, CircuitGNN identified the monosynaptic connection from the interneuron B52 to the interneuron B64, which terminates the retraction phase of feeding behavior. Together with the reciprocal inhibitory connection from B64 to B52, this connection underlies the phasic nature of the feeding behavior [38]. CircuitGNN also identified an inhibitory connection from B52 to motor neuron B8, which regulates the phasing of B8 activity and thereby helps determine the type of feeding behavior. However, CircuitGNN failed to detect some connections (e.g., B63 to B30 and B65 to B20) and inferred some that do not exist (e.g., B8 to B35). Moreover, CircuitGNN inferred the connection from B34 to B63 as inhibitory rather than excitatory, whereas DI recovered it correctly.

To assess the importance of the B63 to B31 and CBI-2 to B63 connections for initiating feeding behavior in the computational model, each connection was removed individually while the remainder of the network was left intact. Removing either connection completely abolished phasic feeding behavior (Fig. 5B, C). Together, these results show that the circuit inferred by CircuitGNN captures several important mechanisms underlying the behavior mediated by the circuit.

**Figure 5.**
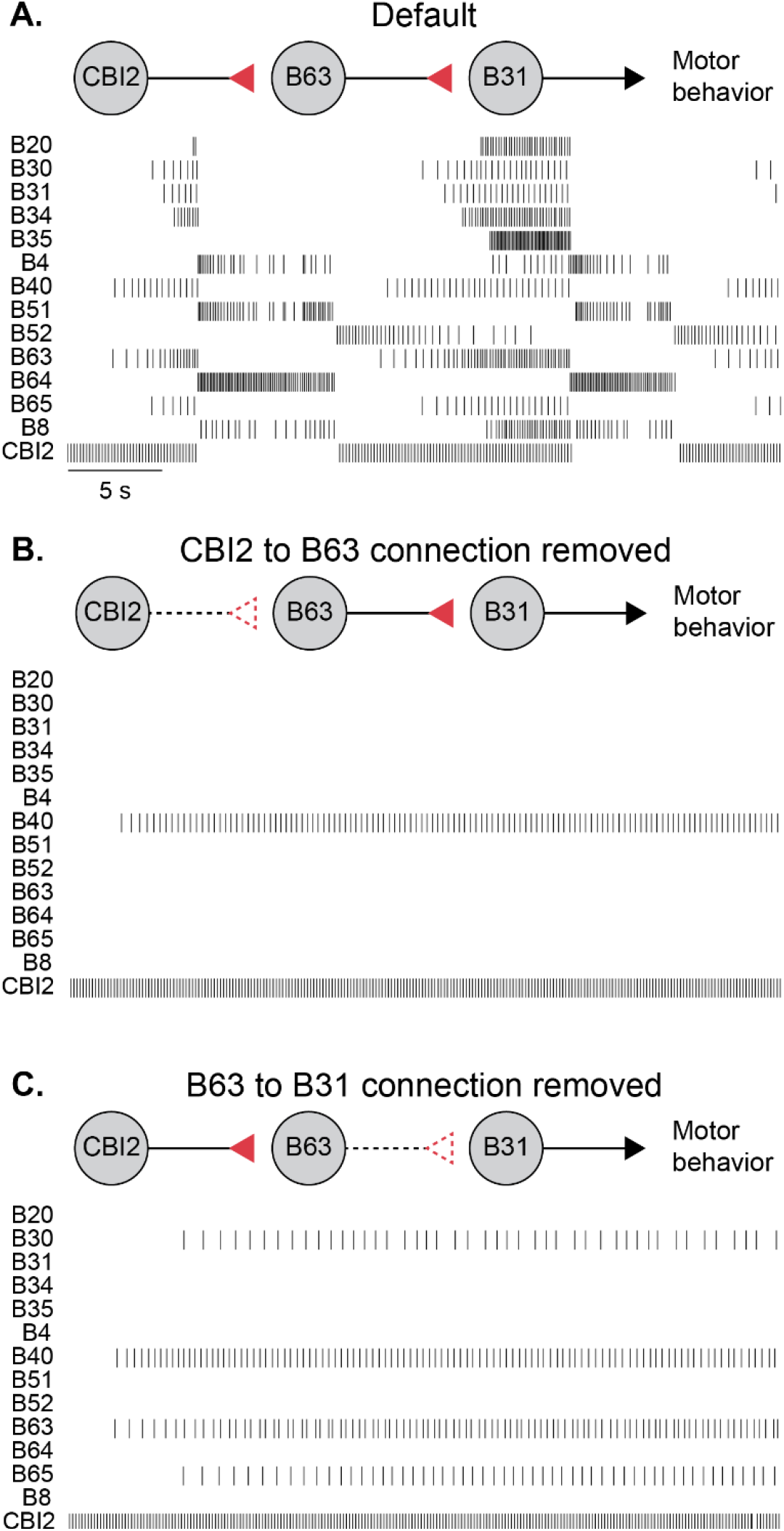
CircuitGNN identified connections central to the behavior generation of the *Aplysia* feeding circuit. **A**. Spike raster of the simulated feeding central pattern generator during operant conditioning; same as Fig. 4A. **B**. Spike raster of the simulated feeding central pattern generator with the excitatory CBI-2 to B63 connection removed. The network is unable to produce rhythmic feeding behavior. **C**. Spike raster of the simulated feeding central pattern generator with the excitatory B63 to B31 connection removed. Similar to B, the network is unable to produce rhythmic feeding behavior.

### CircuitGNN is robust to short recordings and hidden neurons

CircuitGNN was next applied to a simulated cortical network of ~150 neurons [39,40]. The Potjans-Diesmann cortical model (PDCM) is a full-scale model of a layered cortical microcircuit, comprising spiking leaky integrate-and-fire neurons connected by static synapses (i.e., no plasticity). The model includes eight excitatory and inhibitory populations in cortical layers 2/3, 4, 5, and 6, with population sizes, connection probabilities, synaptic strengths, and external inputs constrained by experimental data compiled from a wide range of studies. The PDCM has been used to investigate the emergence of macroscopic cortical patterns [41] and the computational properties of cortical columns [42]. Because the full-scale model contains ~80,000 neurons, it was scaled down to 0.2% of its original size for computational feasibility (Fig. 6A, B). The scaling method adjusts the number of neurons in each population, connections per population, and synaptic weights to preserve the first (mean) and second (variance) order statistics of network activity, thereby retaining layer-specific average firing rates, synchrony, and irregularity features across different network sizes [40].

**Figure 6.**
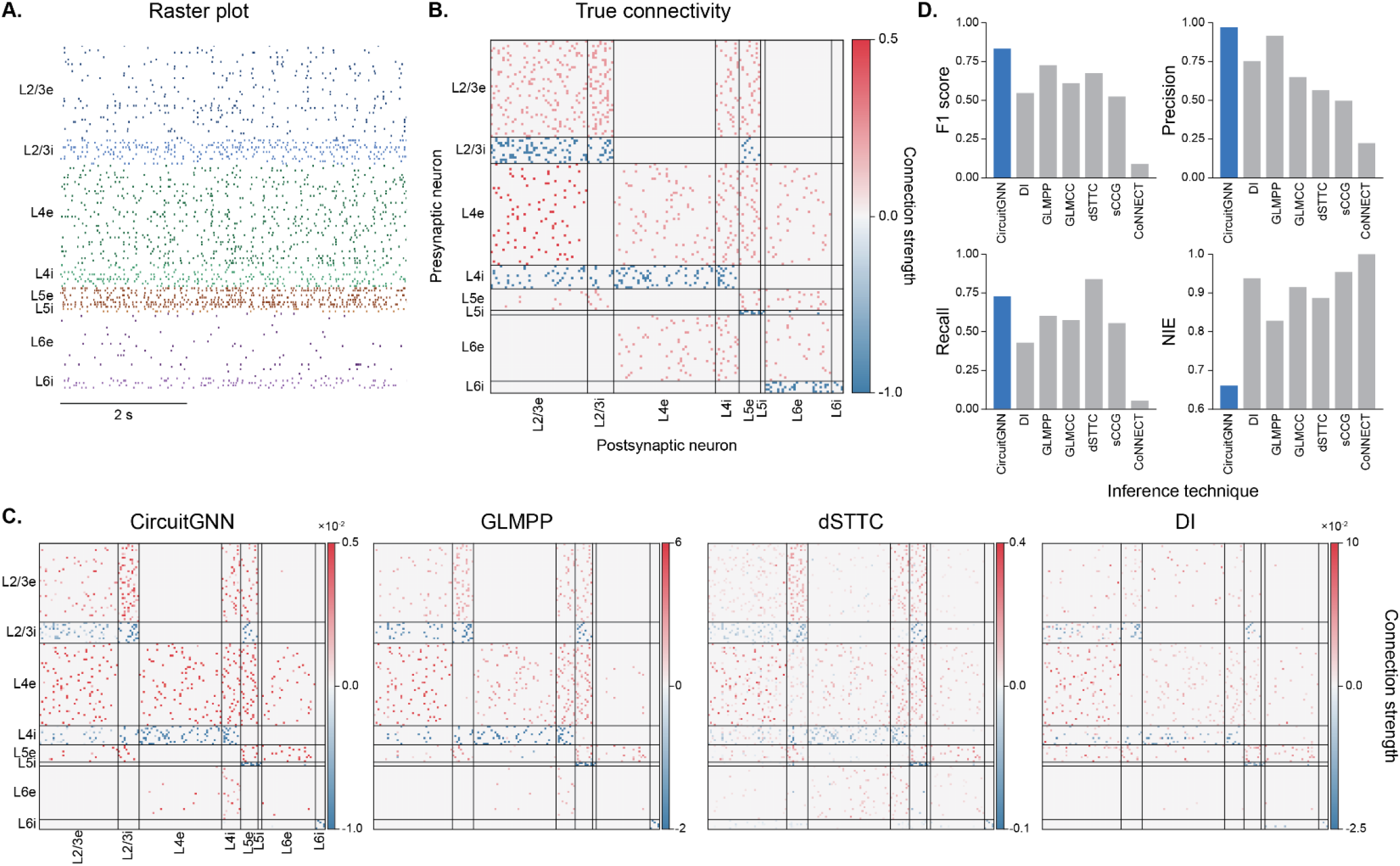
CircuitGNN outperforms existing techniques on inferring the connectivity of a cortical microcircuit. **A**. Spike raster of the simulated Potjans-Diesmann cortical microcircuit, colored by population. **B**. Ground truth connectivity, normalized by the largest synaptic weight. Black lines delimit the eight populations. **C**. Connectivity inferred by CircuitGNN and three of the other techniques. **D**. F1 score, precision, recall, and normalized inference error for all evaluated techniques: CircuitGNN, directed information (DI), generalized linear model for point-processes (GLMPP), generalized linear model cross-correlogram (GLMCC), directed spike time tiling coefficient (dSTTC), smoothed cross-correlogram (sCCG), and convolutional neural network for estimating synaptic connectivity from spike trains (CoNNECT).

CircuitGNN and the other connectivity inference techniques were applied to the PDCM network. CircuitGNN obtained an F1 score of 0.83 and an NIE of 0.66 (Fig. 6C, D, SFig. 3). CircuitGNN’s F1 score was 0.1 higher and its NIE was 0.17 lower than those of the next best technique, and it maintained a high precision@K over more connections than other techniques (SFig. 2B). Together, these results suggest that CircuitGNN outperforms existing methods across multiple metrics. Interestingly, all techniques, including CircuitGNN, failed to identify L6i-to-L6e inhibitory connections (Fig. 6C).

A useful inference model must recover connectivity accurately across diverse recording conditions and circuit sizes. The performance of CircuitGNN was therefore examined as the PDCM network was scaled across two dimensions: number of neurons (149 and 767) and recording duration (60, 120, 240, 480 s). Increasing the size of the network to 767 neurons reduced the performance of CircuitGNN, and increasing recording duration improved the F1 score on both networks (Fig. 7A1). This analysis showed that increasing recording duration improved precision, with values near 0.95 for both networks at recordings as short as 4 min, whereas recall remained relatively unchanged from its value at the shortest recording. Longer recordings also reduced NIE, although the improvement was modest (Fig. 7A2). This result demonstrates two important points: first, because CircuitGNN achieves high precision with only 4 min of recording, many existing datasets that were not originally collected for connectivity inference can be repurposed to infer connectivity; second, CircuitGNN infers connectivity with high precision for circuits as large as ~750 neurons. However, recordings longer than 4–8 min are likely to offer minimal additional benefits, as precision saturates (0.98 for the 149-neuron network, 0.99 for the 767-neuron network at 8 min) and recall remains largely unaffected by duration.

**Figure 7.**
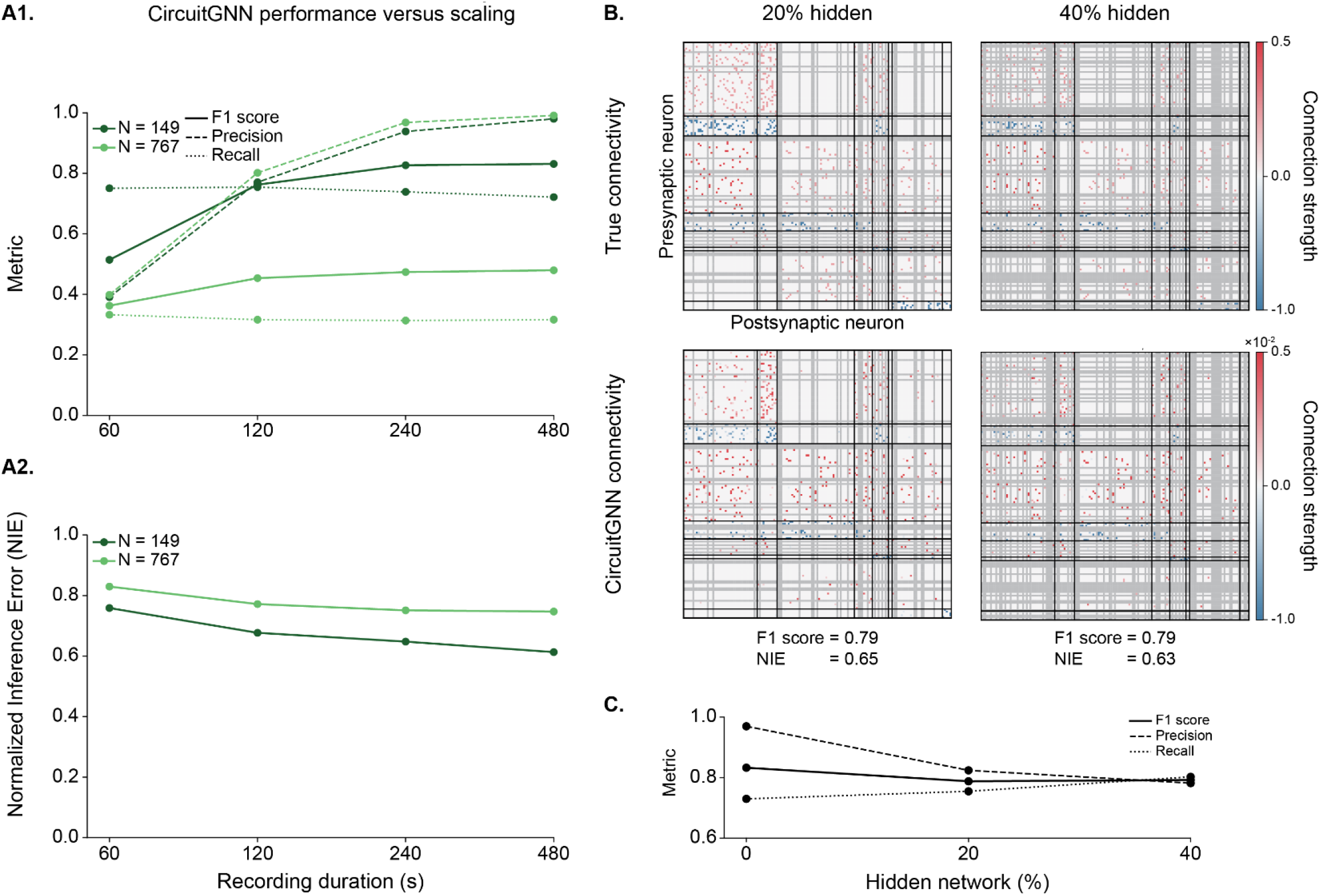
CircuitGNN is robust to short recordings, large networks, and unobserved neurons. **A1**. F1 score, precision, and recall and **A2**. normalized inference error as a function of recording duration, for the 149- and 767-neuron networks. **B**. Ground truth (top) and inferred (bottom) connectivity with 20% (left) and 40% (right) of the 149-neuron network hidden; grey rows and columns mark hidden neurons, which are excluded from all metrics. **C**. F1 score, precision, and recall as a function of the hidden fraction.

A further challenge in connectivity reconstruction is the problem of unobserved neurons (i.e., cells that shape the activity of recorded neurons but are not themselves recorded [34,43]). To test the robustness of CircuitGNN to unobserved neurons, the performance was reassessed after a varying proportion of neurons in the 149-neuron PDCM network (Fig. 6) was hidden. Randomly hiding 20% and 40% of the network yielded an F1 score of 0.79 in both cases, a reduction of just 0.04 relative to the fully observed network (Fig. 7B). As the fraction of hidden neurons increased from 0% to 20% to 40%, precision decreased whereas recall increased, with the two changes largely offsetting one another (Fig. 7C).

Together, these results highlight that CircuitGNN can accurately infer the connectivity of large, cortical networks. Moreover, CircuitGNN achieves its optimal performance at relatively short recording durations and is robust to partially observed networks.

## DISCUSSION

CircuitGNN is a graph structure learning technique, combining a structure learner that infers the underlying connectivity with a GNN that uses this connectivity to optimize a downstream task: forecasting neural activity. CircuitGNN was first tested across several toy networks previously used to benchmark connectivity inference algorithms [9,10]. CircuitGNN accurately recovered the underlying connectivity of these networks (Fig. 2). CircuitGNN also detected changes in a network, such as a reduction or increase in connection strength (Fig. 3). These results demonstrate that CircuitGNN can measure relative strength. This capability allows weak and strong synapses to be distinguished and connection strengths to be compared across experimental conditions, both of which can further elucidate the underlying circuit.

CircuitGNN inferred from population activity the connectivity of computational models of invertebrate (Fig. 4) and cortical (Fig. 6) circuits with high accuracy, outperforming other techniques. In both cases, CircuitGNN maintained higher precision than the other approaches without a significant reduction in recall. Only one technique (dSTTC) achieved higher recall than CircuitGNN, and it did so at substantially lower precision. Thus, whereas CircuitGNN fails to identify some true connections, the large majority of those it does identify are correct. CircuitGNN can therefore offer high-confidence predictions about individual connections and connectivity patterns, narrowing the range of hypotheses about a given circuit.

Several extensions could further strengthen the proposed approach. First, the training objective could be expanded beyond spiking activity. For example, a promising approach may be the pairing of neural recordings with behavioral measures (e.g., head direction, speed) [44]. Second, prior knowledge of the circuit could be incorporated. Incorporating the structural connectome improves estimates of causal relationships between neurons [45], and structural connectomes are available for a growing number of circuits [1–3,46]. For example, for the *Aplysia* CPG circuit, the connections between many pairs of neurons have been characterized [30,31], and individual neurons can be identified in large-scale recordings [47]. The corresponding entries of the adjacency matrix could therefore be initialized with these known connections, allowing for more accurate inference of the remaining connectivity.

Like all connectivity inference techniques, CircuitGNN is correlative, so causal claims about inferred connections should be made with caution, and it can only infer connectivity between neurons that are active during the recording [34,43]. In some cases, such as the *Aplysia* CPG circuit (Fig. 4D) and the 767-neuron cortical network (Fig. 7A1), CircuitGNN detected fewer than half of all connections, so the connectivity it infers likely represents only a subset of all the connections. CircuitGNN was unable to detect inhibitory connections between two populations with relatively low firing rates (Fig. 6, L6i to L6e), suggesting that sparse activity may limit its accuracy. Finally, its performance decreased with increasing network size (Fig. 7A), suggesting that it may not scale well to very large circuits of thousands of neurons. Nevertheless, CircuitGNN outperformed other methods across multiple datasets and performance measures, representing a substantial advance over current approaches.

The advantages of inferring neural connectivity extend beyond characterizing the underlying circuit, to applications such as improved brain-machine interfaces [48] and improved diagnosis of brain disorders. Altered connectivity is a hallmark of several brain disorders, such as Alzheimer’s disease and schizophrenia [49], so accurate inference of brain networks could yield improved diagnostic markers. Reduced or increased connectivity, for example, could be a risk factor or indicator of disease progression. Although CircuitGNN was applied in the present study to infer connectivity at the microscale, its key assumption, that a node’s future activity depends on its own past activity and that of its neighbors, is not specific to individual neurons. The same framework could therefore be applied to infer connectivity at the mesoscale (e.g., from fiber photometry) or the macroscale (e.g., from fMRI), where nodes represent neural populations or brain regions rather than single cells.

## ACKNOWLEDGEMENTS

We thank Arlei Silva and Paul Smolen for their comments on an earlier draft of the manuscript. This work was supported by National Institutes of Health grant R01 NS101356.

## DATA AVAILABILITY STATEMENT

The code for CircuitGNN, including detailed usage instructions and scripts to reproduce the results of this manuscript, is freely available at github.com/Byrne-Lab/circuitGNN. The datasets used in this manuscript are available at doi.org/10.5281/zenodo.23674725.

## METHODS

### CircuitGNN

#### Problem setup

The goal of inferring connectivity between neurons is a graph structure learning (GSL) problem, in which a graph is learned through the optimization of a downstream task. Here, the downstream task being optimized is a forecasting problem: prediction of future spiking activity from past activity. The spiking activity of *N* neurons over *T* discrete time bins is recorded; let 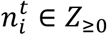 denote the binned spike count of neuron *i* at time *t*, and 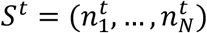 the population state at time *t*. The circuit is modeled as a directed graph *G* = (*V, E*) whose nodes *V* are neurons and edges *E* form a connectome *A* ∈ ℝ^*N*×*N*^, with *A*_*ij*_ quantifying the directed influence of neuron *j* (presynaptic) on neuron *i* (postsynaptic). *A* is inferred jointly with a next-step spike predictor; the whole model is trained in a self-supervised manner based on neuron activity and using no prior knowledge of ground truth connectivity.

#### Spike encoder

To capture short-term dynamics (Fig. 1B), the recent history of each neuron is summarized over a window of length *w*. Let 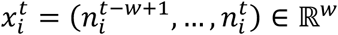. First, each neuron’s window is standardized using its own average spike count:

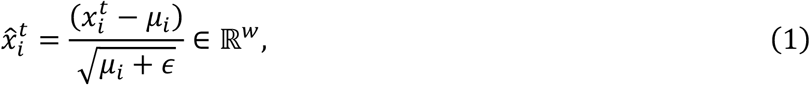

where µ_*i*_ is neuron *i*’s mean spike count per bin computed over the full recording, 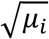 is an estimate of each neuron’s standard deviation, and ϵ is a small constant added for numerical stability. Second, a convolutional encoder maps the standardized window to a feature vector:

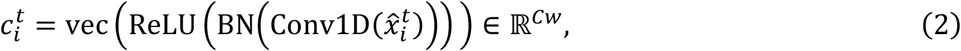

where Conv1D applies *C* learnable filters of kernel size *k* along the time axis of the window, producing *C* feature channels of length *w*. Batch normalization (BN) rescales each channel to zero mean and unit variance across the training batch, which stabilizes training. The rectified linear unit (ReLU) introduces nonlinearity, allowing the encoder to capture nonlinear temporal patterns. Then, vec(·) flattens the resulting *C* × *w* feature map into a single vector 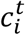 of length *Cw*. Each neuron also carries a persistent memory vector *m*_*i*_ ∈ ℝ^*q*^ that is learned jointly with the rest of the network. Finally, the feature vector 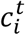 is concatenated with the neuron’s memory vector *m*_*i*_ and passed through a linear projection to produce a low-dimensional embedding:

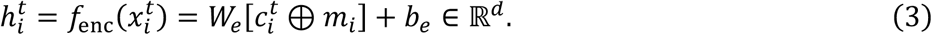

The convolution filters and projection weights (*W*_*e*_, *b*_*e*_) are shared across all neurons whereas *m*_*i*_ is neuron-specific.

#### Connectivity matrix

To distinguish synaptic types, connectivity is parameterized with two unconstrained matrices Θ^+^, Θ^−^ ∈ ℝ^*N*×*N*^ and their rectified parts are taken as nonnegative excitatory and inhibitory components, *A*^*exc*^ = *ReLU*(Θ^+^) ≥ 0 and *A*^*inh*^ = *ReLU*(Θ^−^) ≥ 0. The signed connectome is:

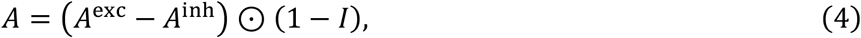

with the diagonal removed to prevent self-edges. This formulation allows for the capture of excitatory and inhibitory connections (Fig. 1C1). These matrices are initialized as the absolute values of a standard normal distribution, scaled by 0.01. The adjacency matrix provides the strength, direction, and type (excitatory/inhibitory) of connections between neurons. For a given connection, the absolute value is the strength and the sign designates whether the connection is excitatory (positive) or inhibitory (negative).

*Message passing*. Neuron representations are refined through *L* layers of message passing that use the connectivity matrix *A* (Fig. 1C2). With 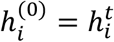, each layer *l* = 1, …, *L* combines three terms. The first is the self-term, which relies solely on each neuron’s own embedding:

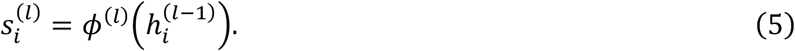

The second is the additive term, which aggregates information from every other neuron, weighted by

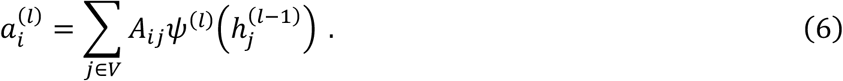

The third is the multiplicative term, which expresses how a neuron’s own state gates the influence of every other neuron, also weighted by *A*:

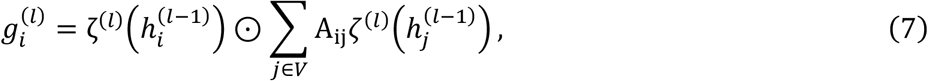

here ⊙ represents the Hadamard product. These three terms are combined using learned scalars *β*_1_, *β*_2_, *β*_3_ that weight their relative contributions (Fig. 1C3). The final representation of neuron *i* in layer *l* is:

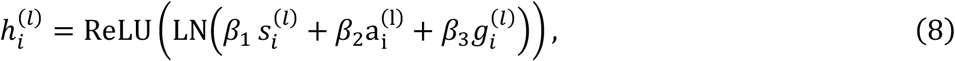

where *ψ*^(*l*)^ and *ζ*^(*l*)^ are two-layer MLPs, *ϕ*^(*l*)^ is a linear layer with ReLU, and LN is layer normalization. By reusing *A* across layers, its influence propagates over multi-hop paths while still yielding a single inferred connectome.

*Decoder and objective*. A linear decoder maps the final representation of each neuron to a scalar logit:

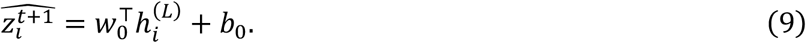

Passing this logit through a sigmoid gives the predicted probability that neuron *i* spikes in the next bin (Fig. 1D):

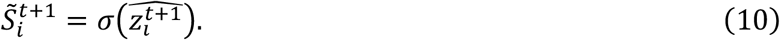

The target is whether neuron *i* spikes in the following bin, 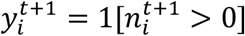. Because spikes are sparse, a class-weighted binary cross-entropy is optimized over *T*′ prediction windows (*T*^′^ < *T* because the first *w* bins of the recording do not form a complete input window):

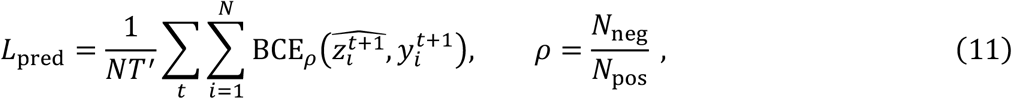

where the positive-class weight ρ is estimated from the training split. The first 80% of time bins are used for training, and the remaining 20% for validation.

#### Connectivity regularization

Two penalties regularize the inferred connectivity *A*. A log-magnitude penalty promotes sparsity:

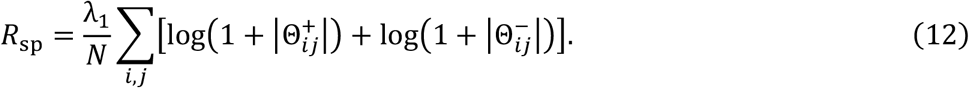

A competition penalty discourages any single edge from being simultaneously excitatory and inhibitory, driving each connection toward a definite sign:

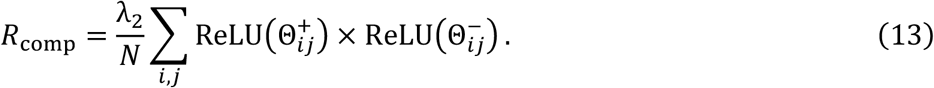

Both terms are normalized per neuron. The full objective is *L* = *L*_pred_ + *R*_sp_ + *R*_comp_, minimized with Adam; the checkpoint with the lowest validation cross-entropy *L*_pred_ is retained.

#### Replicates

For each dataset, 50 replicates were trained with random initializations. For the scaling and hidden-neuron analyses (Fig. 7), 20 replicates were trained for each combination of network size and recording duration and for each proportion of hidden neurons. Replicates were first screened by an ablation test. For each replicate, the validation loss of the full model was compared with that of the same model with the learned adjacency zeroed, and any replicate for which ablation increased the loss by less than 0.10 was discarded, indicating that the inferred graph contributed negligibly to prediction. The remaining replicates were sign-oriented (see below) and averaged. The averaged weights were rounded to three decimal places. To suppress noisy edges, only edges whose sign agreed across at least 80% of replicates were retained, and all other edges were set to zero, yielding the final connectivity matrix. The 80% threshold was chosen to balance the tradeoff between precision and recall. Examples of individual replicates are shown in SFig. 4.

#### Sign orientation

Because training provides no absolute sign reference, independent runs can converge to sign-flipped versions of the same connectivity – positive values may denote excitatory connections in one replicate and inhibitory connections in another. This sign ambiguity is resolved in two steps. First, each replicate is aligned to the first: a replicate is flipped whenever its sign-flipped adjacency has higher cosine similarity to the reference. This alignment brings the replicates into agreement with one another but leaves the overall orientation of the set undetermined. Second, a model-free sign estimate is derived from the spike data binned at 20 ms, using the partial-correlation sign inference of [10]. Each edge takes the sign of its partial correlation at the lag of largest magnitude. The entire aligned set is flipped if the summed product of its mean weights with these reference signs is negative. This estimate serves only to resolve the global sign.

#### Bin size selection

CircuitGNN’s temporal resolution is set by the bin width used to build the population state, which is selected heuristically as the timescale at which directed coupling is most reproducible. For each candidate width (1–10 and 20 ms), spikes are binned, up to 150 neurons are sampled, and the recording is divided into four consecutive, non-overlapping segments. Within each segment, a lag-1 vector autoregression is fit, predicting each neuron’s activity from that of all neurons one bin earlier, yielding a coefficient for each pair. Each connection is labeled as excitatory, inhibitory, or absent from the sign and size of its coefficient, using a cutoff of 1 SD. The stability of these inferred labels across segments is quantified at each width as the mean pairwise Cohen’s *κ* [50], and the width that maximizes this value is selected. If Cohen’s *κ* rises with bin width rather than peaking within the tested range, a default of 4 ms is used. Bin sizes of 9 ms, 4 ms, and 3 ms were chosen for networks A, B, and the PDCM network, respectively, whereas the 4 ms default was used for network C and the *Aplysia* network. For the scaling and hidden neuron PDCM variants (Fig. 7), the bin width was set to 3 ms to match the PDCM network.

#### CircuitGNN parameters

Most hyperparameters (e.g., embedding dimension, message MLP hidden width) were set to fixed values without tuning (Table 1). For a few others (e.g., sparsity weight, competition weight), several values were tested and those yielding accurate connectivity inference were selected. In these tests, CircuitGNN’s performance was largely insensitive to moderate changes in most hyperparameters. The same hyperparameter values were used for all datasets.

**Table 1.** CircuitGNN hyperparameters.

| Component | Parameter | Value |
|---|---|---|
| Input | Window length (bins) | 100 |
| Encoder | Convolutional filters | 32 |
|  | Kernel size | 3 |
|  | Memory vector dimension | 32 |
|  | Embedding dimension | 64 |
| GNN | Message-passing layers | 5 |
|  | Hidden representation size | 64 |
|  | Message MLP hidden width | 128 |
|  | Output dimension | 1 |
| Training | Optimizer | Adam |
|  | Learning rate | 10<sup>−4</sup> |
|  | Batch size | 16 |
|  | Epochs | 20–50 |
|  | Train/validation split | 80/20 |
| Regularization | Sparsity weight | 2.0 |
|  | Competition weight | 15.0 |

### Benchmark techniques

#### Smoothed cross-correlogram (sCCG)

Given two spike trains, a cross-correlogram (CCG) quantifies the number of coincident spikes at the same instant (zero lag) and across a range of time lags. Interpreting the CCG requires testing it against the null hypothesis that the two trains are uncorrelated at a precise temporal resolution. Therefore, a predictor cross-correlogram was computed by convolving the raw CCG with a partially hollow Gaussian kernel, in which the zero lag bin was down-weighted [35]. sCCG was computed with the *spycon* Python package [17]. Bin size and hollow fraction were varied jointly, over all combinations of bin size = [1 ms, 2 ms, 5 ms] and hollow fraction = [0.4, 0.6, 0.8]. Three additional parameters were each varied individually: the synaptic window over [0.5, 4], [1, 10], and [2, 20] ms; the significance level α over 0.01 and 0.05; and CCG deconvolution, enabled and disabled. All parameters not being varied were set to their default values. The best-performing setting was a bin size of 2 ms with a hollow fraction of 0.8 for the PDCM network, and a significance level of 0.05, with all other parameters at their default values, for the *Aplysia* CPG network.

#### Directed spike time tiling coefficient (dSTTC)

dSTTC is a directed variant of the spike time tiling coefficient (STTC), a measure of temporal overlap between two spike trains within a synchrony window [36]. Whereas STTC symmetrically averages both directions, dSTTC yields a directed estimate [17]. dSTTC was computed with the *spycon* Python package [17]. Four parameters were each varied individually: the synchrony window over 3, 5, 7, 10, 20 and 30 ms; the number of surrogates over 20 and 100; the significance level over 0.01 and 0.05; and a jitter-based null, whose jitter factor was varied over 3, 7, and 15. All parameters not being varied were set to their default values. The best-performing setting was synchrony window = 30 ms for both the PDCM and *Aplysia* CPG networks.

#### Generalized linear model for point-processes (GLMPP)

GLMPP is a point-process generalized linear model that predicts each neuron’s firing rate from the recent spiking history of the entire circuit [37]. The fitted cross-neuron coupling terms provide a directed estimate of connectivity. Such models have previously been used to infer connectivity [51]. GLMPP was computed with the *spycon* Python package [17] with all parameters at their default values.

#### Generalized linear model cross-correlogram (GLMCC)

GLMCC infers connectivity by fitting a generalized linear model to the pairwise spike train cross-correlogram, in which a slowly varying background term and fast synaptic coupling filters contribute additively to the firing rate. The magnitude of the fitted coupling parameter, expressed in units of postsynaptic potential, measures connection strength [13]. GLMCC was computed with the *spycon* Python package [17]. The synaptic delay, synaptic decay time constant, and the regularization weight were varied jointly, over all combinations of synaptic delay = [1 ms, 3 ms, 5 ms, 10 ms], synaptic decay time constant = [1 ms, 4 ms], and regularization weight = [500, 4000]. Two additional parameters were varied jointly at a synaptic delay of 10 ms: the cross-correlogram time constant over 25 and 100 ms, and CCG deconvolution, enabled and disabled. All parameters not being varied were set to their default values. The best-performing parameters were a synaptic delay of 1 ms with synaptic decay time constant = 4 ms and regularization weight = 4000 for the PDCM network, and a synaptic delay of 5 ms with synaptic decay time constant = 4 ms and regularization weight = 500 for the *Aplysia* CPG network.

#### Directed information (DI)

DI is an information theoretic measure that quantifies the directed exchange of information between two systems and has previously been applied to infer connectivity [9,10]. DI was calculated using the CTM-DI method [9]. Because this implementation automatically evaluates DI across bin sizes from 2 to 30 ms in 1 ms increments, no parameters needed to be manually tuned to optimize performance. Following the published approach [9], a connection was identified only if its DI value exceeded 0.01 for four consecutive bin sizes, and all nonzero values were then averaged to produce the final estimate.

#### CoNNECT

Convolutional Neural Network for Estimating synaptic Connectivity from spike Trains (CoNNECT) is a functional connectivity inference method developed by Endo et al. [16]. A convolutional neural network was trained on thousands of cross-correlogram images to detect the presence of a monosynaptic connection between a pair of neurons and to estimate the amplitude of the postsynaptic potential that one neuron would drive in another. The method includes no user-adjustable parameters. Spike times were binned at 1 ms resolution prior to analysis.

### Performance quantification

#### F1 score

F1 score is a measure of a classifier’s predictive performance. Each possible connection is labeled as excitatory, inhibitory, or no connection. An inferred connection counted as a true positive (TP) only if the ground truth connection existed between the same neurons with the same sign. An inferred connection that was either absent in the ground truth or present with the opposite sign counted as a false positive (FP). A ground truth connection that was left uninferred or inferred with the opposite sign counted as a false negative (FN). Precision is 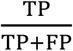, recall is 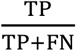, and F1 score is their harmonic mean, 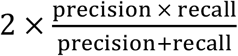. Counts were pooled across the excitatory and inhibitory classes before calculating these metrics. Higher values indicate more accurate performance.

#### Normalized inference error

Normalized inference error (NIE) is a continuous measure of how closely the inferred weights match the ground truth [25,34]. Connection weights inferred by CircuitGNN may differ from the true weights by an arbitrary global scale. To eliminate the effect of such scaling, a single scalar is chosen that minimizes the *l*_1_-distance between the inferred and true connectivity matrices.

Then, the quality of the inferred matrices is calculated as 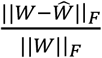, where *W* is the true connectivity, 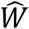 is the inferred connectivity, and ||. ||_*F*_ denotes the Frobenius norm. Lower values indicate more accurate performance.

### Datasets

#### Small circuits

Networks A, B, and C are conductance-based models simulated in SNNAP [29], yielding membrane potential values at 0.5 ms (A, B) or 1 ms resolution (C). Spikes were extracted by thresholding the membrane potential at −10 mV with a re-arm period of 8 ms to count one spike per action potential. As was done in [9] and [10], networks A and B were simulated for 200 s, whereas network C was simulated for 300 s. Ground truth connectivity for network C was obtained from the SNNAP model’s synaptic conductances and normalized by the largest magnitude.

#### Cortical microcircuit

The Potjans-Diesmann model [39] was simulated in NetPyNE [40] at 0.2% (149 neurons) and 1% (767 neurons) of full scale, set via the *cfg*.*ScaleFactor* parameter (0.002 and 0.010, respectively). Spike times and ground truth connectivity were obtained from the simulator. Simulation duration was set via the *cfg*.*duration* parameter: the 149-neuron dataset used for Fig. 6 was simulated for 200 s, and the scaling datasets were simulated for 2000 s and truncated to varying durations (Fig. 7A). All other parameters were kept at their default values.

#### Aplysia feeding circuit

Fourteen neurons were simulated in SNNAP for 240 s at 0.5 ms resolution using the operant conditioning model [31, OCx1 SNNAP simulation] (GitHub: github.com/Byrne-Lab/momohara_neveu_2021/tree/main/OCx1). Spikes were extracted by thresholding the membrane potential at −10 mV with a re-arm period of 8 ms to count one spike per action potential. Ground truth connectivity was obtained from a separate stimulation protocol. Each neuron was depolarized in turn for 20 ms and the response of every other neuron was measured as its mean membrane potential over the following 200 ms minus its membrane potential at stimulus onset. Responses were retained only for pairs connected via chemical and/or electrical synapses.

## SUPPLEMENTARY FIGURES

**SFig. 1.**
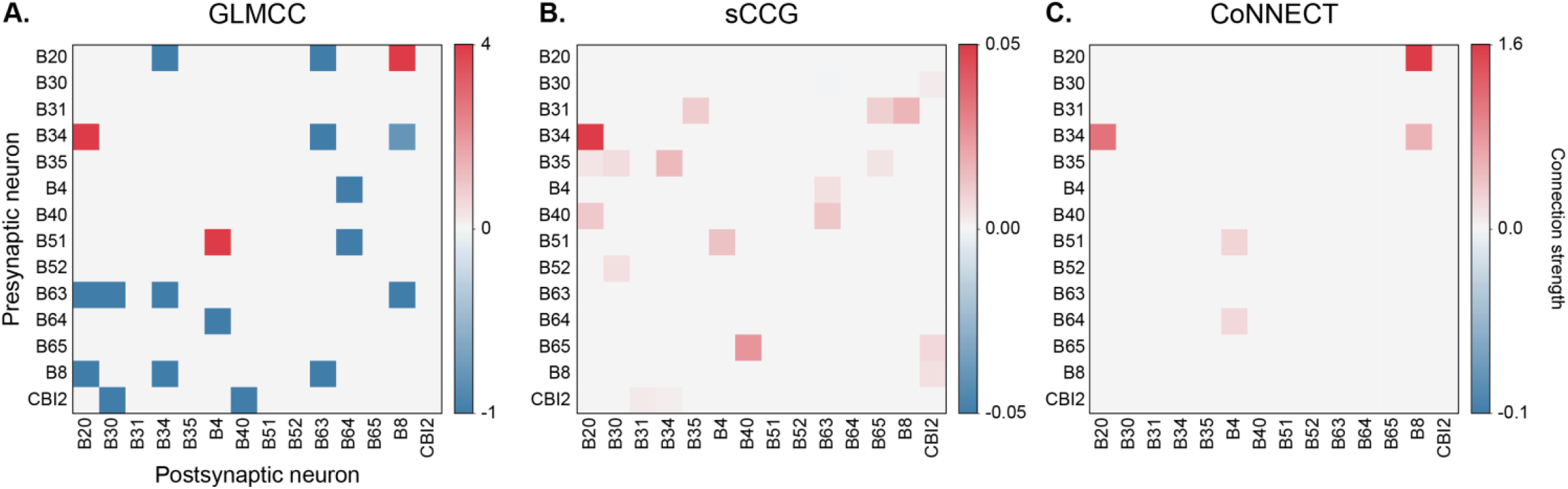
Connectivity matrices of the *Aplysia* feeding circuit inferred by **A**. GLMCC, **B**. sCCG, and **C**. CoNNECT.

**SFig. 2.**
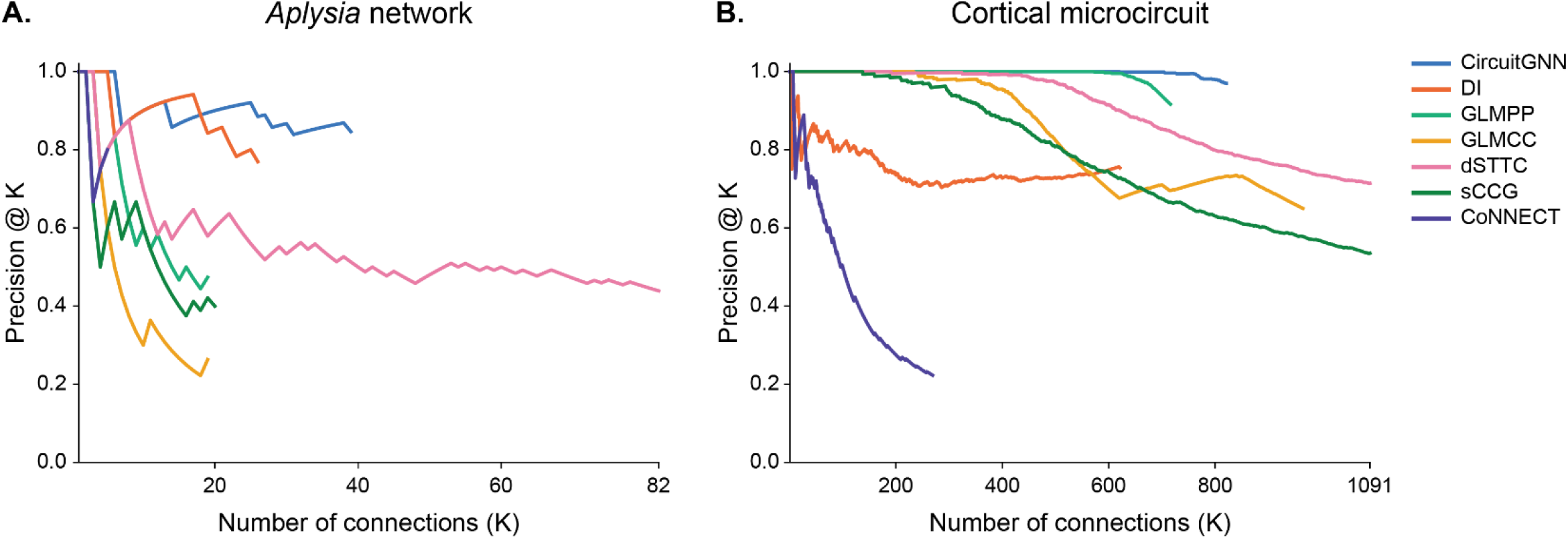
Precision@K of all connectivity inference techniques for **A**. the *Aplysia* network and **B**. the cortical microcircuit. For each technique, predicted connections were ranked by estimated connection strength, and precision@K was computed as the fraction of the K highest-ranked connections that are true connections. K was varied from 1 to the number of connections predicted by each technique, up to the total number of true connections. Therefore, curves for different techniques end at different values of K.

**SFig. 3.**
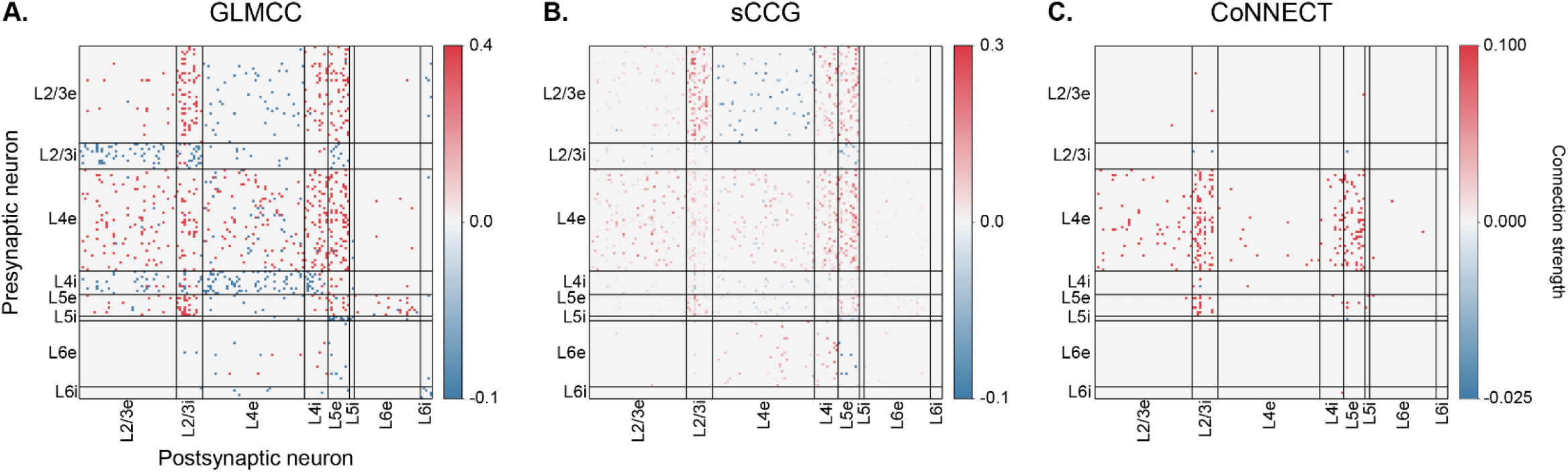
Connectivity matrices of the cortical microcircuit inferred by **A**. GLMCC, **B**. sCCG, and **C**. CoNNECT

**SFig. 4.**
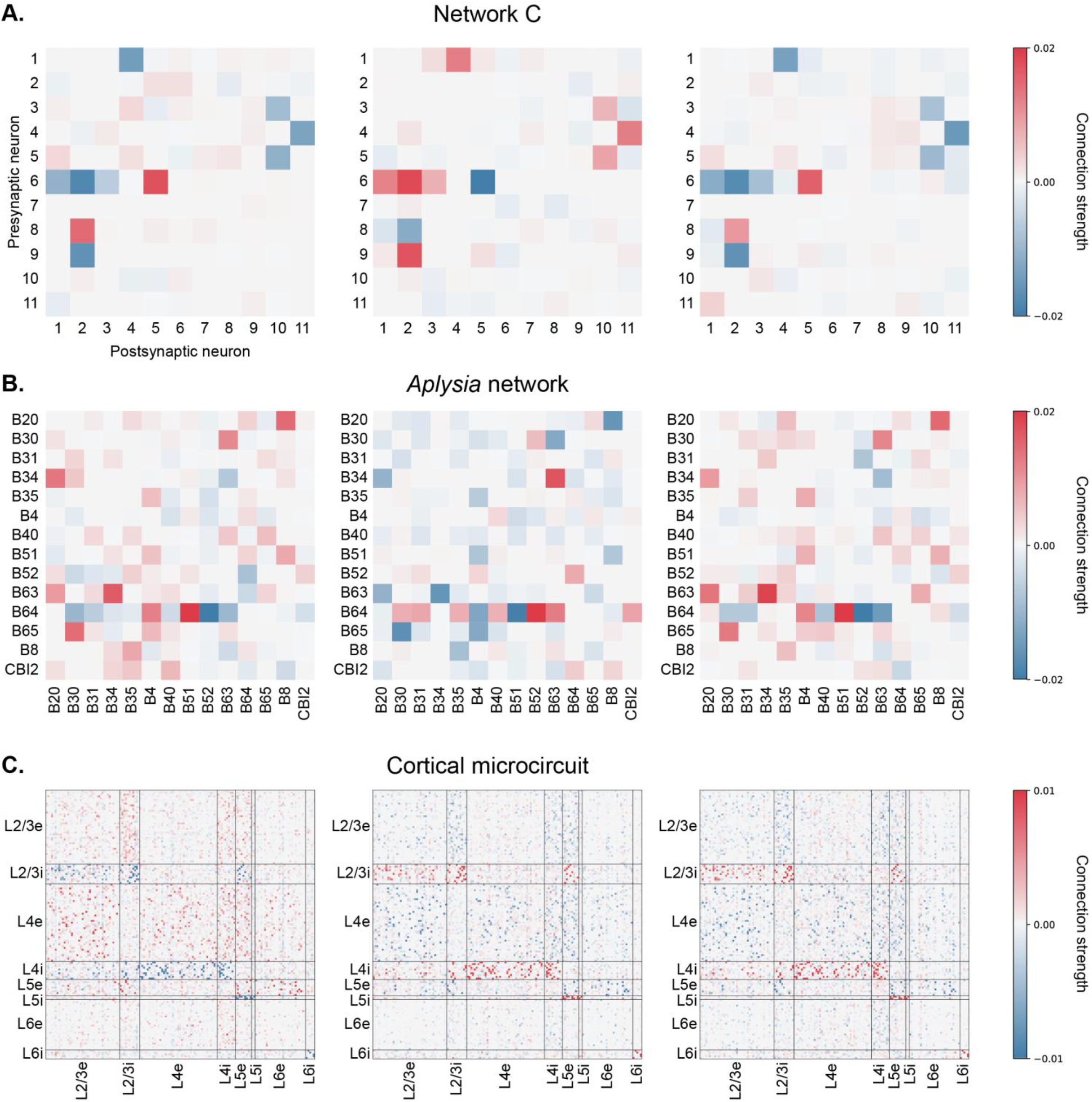
Connectivity inferred by randomly chosen individual CircuitGNN replicates (random initializations) for **A**. Network C, **B**. the *Aplysia* network, and **C**. the cortical microcircuit. The final connectivity matrices reported in the main text combine many replicates per dataset (see Methods).

## REFERENCES

[1] Cook S J et al. 2019 Whole-animal connectomes of both Caenorhabditis elegans sexes Nature 571 63–71

[2] Bae J A et al. 2025 Functional connectomics spanning multiple areas of mouse visual cortex Nature 640 435–47

[3] Berg S et al 2026 Sexual dimorphism in the complete Drosophila male central nervous system connectome Cell 189 5504–5526.e15

[4] Oh S W et al. 2014 A mesoscale connectome of the mouse brain Nature 508 207–14

[5] Mozzachiodi R and Byrne J H 2010 More than synaptic plasticity: role of nonsynaptic plasticity in learning and memory Trends in Neurosciences 33 17–26

[6] Pastore V P, Massobrio P, Godjoski A and Martinoia S 2018 Identification of excitatory-inhibitory links and network topology in large-scale neuronal assemblies from multi-electrode recordings PLoS Comput Biol 14 e1006381

[7] De Blasi S, Ciba M, Bahmer A and Thielemann C 2019 Total spiking probability edges: a cross-correlation based method for effective connectivity estimation of cortical spiking neurons J Neurosci Methods 312 169–81

[8] Ito S, Hansen M E, Heiland R, Lumsdaine A, Litke A M and Beggs J M 2011 Extending transfer entropy improves identification of effective connectivity in a spiking cortical network model PLoS One 6 e27431

[9] Cai Z, Neveu C L, Baxter D A, Byrne J H and Aazhang B 2017 Inferring neuronal network functional connectivity with directed information J Neurophysiol 118 1055–69

[10] Young J, Neveu C L, Byrne J H and Aazhang B 2021 Inferring functional connectivity through graphical directed information J Neural Eng 18 046019

[11] Ekhlasi A, Nasrabadi A M and Mohammadi M 2023 Improving transfer entropy and partial transfer entropy for relative detection of effective connectivity strength between time series Commun Nonlinear Sci Numer Simul 126 107449

[12] Gerhard F, Kispersky T, Gutierrez G J, Marder E, Kramer M and Eden U 2013 Successful reconstruction of a physiological circuit with known connectivity from spiking activity alone PLoS Comput Biol 9 e1003138

[13] Kobayashi R, Kurita S, Kurth A, Kitano K, Mizuseki K, Diesmann M, Richmond B J and Shinomoto S 2019 Reconstructing neuronal circuitry from parallel spike trains Nat Commun 10 4468

[14] Shinomoto S and Tsubo Y 2026 Comparing effective and functional connectivity Sci Rep 16 12161

[15] Pozo-Jimenez P, Lucas-Romero J and Lopez-Garcia J A 2021 Discovering effective connectivity in neural circuits: analysis based on machine learning methodology Front Neuroinform 15 561012

[16] Endo D, Kobayashi R, Bartolo R, Averbeck B B, Sugase-Miyamoto Y, Hayashi K, Kawano K, Richmond B J and Shinomoto S 2021 A convolutional neural network for estimating synaptic connectivity from spike trains Sci Rep 11 12087

[17] Donner C, Bartram J, Hornauer P, Kim T, Roqueiro D, Hierlemann A, Obozinski G and Schröter M 2024 Ensemble learning and ground-truth validation of synaptic connectivity inferred from spike trains PLoS Comput Biol 20 e1011964

[18] Lu Z, Zhang W, Le T, Wang H, Sümbül U, Shea-Brown E and Mi L 2025 NetFormer: an interpretable model for recovering dynamical connectivity in neuronal population dynamics International Conference on Learning Representations

[19] Luo Z, Peng K, Liang Z, Cai S, Xu C, Li D, Hu Y, Zhou C and Liu Q 2025 Mapping effective connectivity by virtually perturbing a surrogate brain Nat Methods 22 1376–85

[20] Magrans de Abril I, Yoshimoto J and Doya K 2018 Connectivity inference from neural recording data: challenges, mathematical bases and research directions Neural Netw 102 120–37

[21] Loomba S et al 2022 Connectomic comparison of mouse and human cortex Science 377 eabo0924

[22] Scarselli F, Gori M, Tsoi A C, Hagenbuchner M and Monfardini G 2009 The graph neural network model IEEE Trans Neural Netw 20 61–80

[23] Corso G, Stark H, Jegelka S, Jaakkola T and Barzilay R 2024 Graph neural networks Nat Rev Methods Primers 4 17

[24] Li J, Scholl L, Le T, Rajeswaran P, Orsborn A and Shlizerman E 2023 AMAG: additive, multiplicative and adaptive graph neural network for forecasting neuron activity Adv Neural Inf Process Syst 36 8988–9014

[25] Yoon K 2025 Self-supervised discovery of neural circuits in spatially patterned neural responses with graph neural networks Adv Neural Inf Process Syst 38 160740–67

[26] Zhu Y, Xu W, Zhang J, Du Y, Zhang J, Liu Q, Yang C and Wu S 2022 A survey on graph structure learning: progress and opportunities arXiv 2103.03036

[27] Mattos J, Lina D H and Silva A 2026 Breaking the dyadic barrier: rethinking fairness in link prediction beyond demographic parity Proc AAAI Conf Artif Intell 40 24335–43

[28] Zhou P, Yin K, Zhu H, Liang Y, Li L and Chen Y 2026 A survey on graph structure learning Neurocomputing 696 134114

[29] Ziv I, Baxter D A and Byrne J H 1994 Simulator for neural networks and action potentials: description and application J Neurophysiol 71 294–308

[30] Costa R M, Baxter D A and Byrne J H 2020 Computational model of the distributed representation of operant reward memory: combinatoric engagement of intrinsic and synaptic plasticity mechanisms Learn Mem 27 236–49

[31] Momohara Y, Neveu C L, Chen H-M, Baxter D A and Byrne J H 2022 Specific plasticity loci and their synergism mediate operant conditioning J Neurosci 42 1211–23

[32] Dickman U, Thomas P J, Chiel H J, Byrne J H and Neveu C L 2026 NEURONpyxl: fast, flexible, Python-integrated simulation of biophysical neural networks with complex plastic synapses Front Comput Neurosci 20 1771884

[33] Christen P, Hand D J and Kirielle N 2023 A review of the F-measure: its history, properties, criticism, and alternatives ACM Comput Surv 56 73

[34] Das A and Fiete I R 2020 Systematic errors in connectivity inferred from activity in strongly recurrent networks Nat Neurosci 23 1286–96

[35] Stark E and Abeles M 2009 Unbiased estimation of precise temporal correlations between spike trains J Neurosci Methods 179 90–100

[36] Cutts C S and Eglen S J 2014 Detecting pairwise correlations in spike trains: an objective comparison of methods and application to the study of retinal waves J Neurosci 34 14288–303

[37] Truccolo W, Eden U T, Fellows M R, Donoghue J P and Brown E N 2005 A point process framework for relating neural spiking activity to spiking history, neural ensemble, and extrinsic covariate effects J Neurophysiol 93 1074–89

[38] Cropper E C, Evans C G, Hurwitz I, Jing J, Proekt A, Romero A and Rosen S C 2004 Feeding neural networks in the mollusc Aplysia Neurosignals 13 70–86

[39] Potjans T C and Diesmann M 2014 The cell-type specific cortical microcircuit: relating structure and activity in a full-scale spiking network model Cereb Cortex 24 785–806

[40] Romaro C, Najman F A, Lytton W W, Roque A C and Dura-Bernal S 2021 NetPyNE implementation and scaling of the Potjans-Diesmann cortical microcircuit model Neural Comput 33 1993–2032

[41] Bos H, Diesmann M and Helias M 2016 Identifying anatomical origins of coexisting oscillations in the cortical microcircuit PLoS Comput Biol 12 e1005132

[42] Cain N, Iyer R, Koch C and Mihalas S 2016 The computational properties of a simplified cortical column model PLoS Comput Biol 12 e1005045

[43] Mehler D M A and Kording K P 2020 The lure of misleading causal statements in functional connectivity research arXiv 1812.03363

[44] Delavari P, Oruc I and Murphy T H 2024 SynapsNet: enhancing neuronal population dynamics modeling via learning functional connectivity arXiv 2411.08221

[45] Pospisil D A et al. 2024 The fly connectome reveals a path to the effectome Nature 634 201–9

[46] Shapson-Coe A et al 2024 A petavoxel fragment of human cerebral cortex reconstructed at nanoscale resolution Science 384 eadk4858

[47] Gupta A, Neveu C L, Cropper E C and Byrne J H 2026 Motor priming is associated with widespread recruitment into neural ensembles and more rapid ensemble transitions iScience 29 117375

[48] Stevenson I H, Rebesco J M, Miller L E and Kording K P 2008 Inferring functional connections between neurons Curr Opin Neurobiol 18 582–8

[49] Bullmore E and Sporns O 2009 Complex brain networks: graph theoretical analysis of structural and functional systems Nat Rev Neurosci 10 186–98

[50] Cohen J 1960 A Coefficient of Agreement for Nominal Scales Educational and Psychological Measurement 20 37–46

[51] Pillow J W, Shlens J, Paninski L, Sher A, Litke A M, Chichilnisky E J and Simoncelli E P 2008 Spatio-temporal correlations and visual signalling in a complete neuronal population Nature 454 995–9

